# The environmental impacts of palm oil and its alternatives

**DOI:** 10.1101/2020.02.16.951301

**Authors:** Robert M. Beyer, América P. Durán, Tim T. Rademacher, Philip Martin, Catherine Tayleur, Sharon E. Brooks, David Coomes, Paul F. Donald, Fiona J. Sanderson

**Author notes:** Authors contributed equally (América P. Durán, Tim T. Rademacher).

## Abstract

The destruction of ecosystems for vegetable oil production represents a major cause of global biodiversity loss and greenhouse gas emissions ^1^. Over the last two decades, oil palm, in particular, has caused societal concern due to its high impacts on biodiverse and carbon-dense tropical rainforests ^2–8^, leading to calls to source vegetable oils from alternative oil-producing crops. However, given the high yields of oil palm, how does that damage compare with other oil crops that require more land? Here, we estimate the carbon and biodiversity footprints, per unit of oil produced, of the world’s five major vegetable oil crops. We find that oil palm has the lowest carbon loss and species richness loss per-tonne-oil, but has a larger impact on range-restricted species than sunflower and rapeseed. We go on to identify global areas for oil crop expansion that will minimise future carbon and biodiversity impacts, and argue that closing current yield gaps and optimising the location of future growing areas will be much more effective at reducing future environmental impacts of global vegetable oil production than substituting any one crop for another.

---

Vegetable oils are among the world’s most rapidly expanding crop types ^9^, and are a major contributor to global biodiversity loss and greenhouse gas emissions caused by land conversion ^1^. Assuming a business-as-usual scenario in which global human population and per capita consumption continue to increase, worldwide demand for vegetable oil may increase from its current level of 205 million Mg ^10^ to up to 340 million Mg by 2050 ^11^. Palm oil, which currently accounts for 41% of global vegetable oil production ^12^, has been repeatedly criticised for its high environmental impacts due to large-scale replacement of tropical forests, resulting in high carbon emissions ^6–8^ and biodiversity loss ^2–5^. Concerns are exacerbated by the ongoing expansion of palm oil production in Southeast Asia, Sub-Saharan Africa and South America ^3,13^. Vegetable oils from different oil crops are often substitutable in terms of end use ^14^, which has led to suggestions that the environmental damage of oil production could be reduced by sourcing from crops other than oil palm. However, whilst the impact of oil palm per hectare is large, its high yield levels of up to ten times those of its alternatives ^15^ mean that relatively small areas need to be cropped to produce a given quantity of oil. This may potentially result in a smaller total environmental impact compared to lower yielding crops ^16^.

Here, we use a spatially explicit framework to map three environmental impacts – carbon loss, species richness loss and range rarity loss – in relation to global yield maps of oil sourced from oil palm, soybean, rapeseed, sunflower and groundnut. Between them, these crops account for 94% of the current global vegetable oil production ^12^. We consider both existing and potential growing areas of each crop, which allows us to rank present as well as potential future growing locations according to their relative environmental impacts (i.e. the ratio of local carbon and biodiversity loss associated with cultivation to local oil yield), and provide recommendations to minimise the footprint of global vegetable oil production.

We used global maps of harvested areas and yields for the year 2010 ^17^, the most recent available spatial data (Methods). Environmental impacts were mapped as follows: Carbon loss was calculated as the difference between local above- and below-ground carbon stocks of (i) natural land cover and (ii) oil cropland ^18^ (Methods). Species richness loss was estimated by overlaying global range maps of all known birds, mammals and amphibians, and determining species lost locally when natural habitat is converted to cropland ^19^ (Methods). Range rarity loss, a biodiversity metric alternative to species richness loss ^20^, is based on the same approach, but species with narrow geographical ranges were weighted more heavily (Methods).

As would be expected from the location of current cropping areas, absolute carbon and biodiversity impacts per hectare are, on average, higher for oil palm than for the other four crops (Fig. S1). We find, however, that this is no longer the case when taking into account the different area requirements of each crop for producing a given quantity of oil. On current growing areas, oil palm has the lowest average carbon loss per-tonne-oil (Fig. 1A). It also has the lowest average species richness loss per-tonne-oil (Fig. 1B), but has a larger impact on range rarity than sunflower and rapeseed (Fig. 1C). The spatial variability in impact per-tonne-oil is substantial for all crops, indicating that growing location affects environmental impacts of vegetable oil production more than crop type. We return to this point later on.

**Fig. 1:**
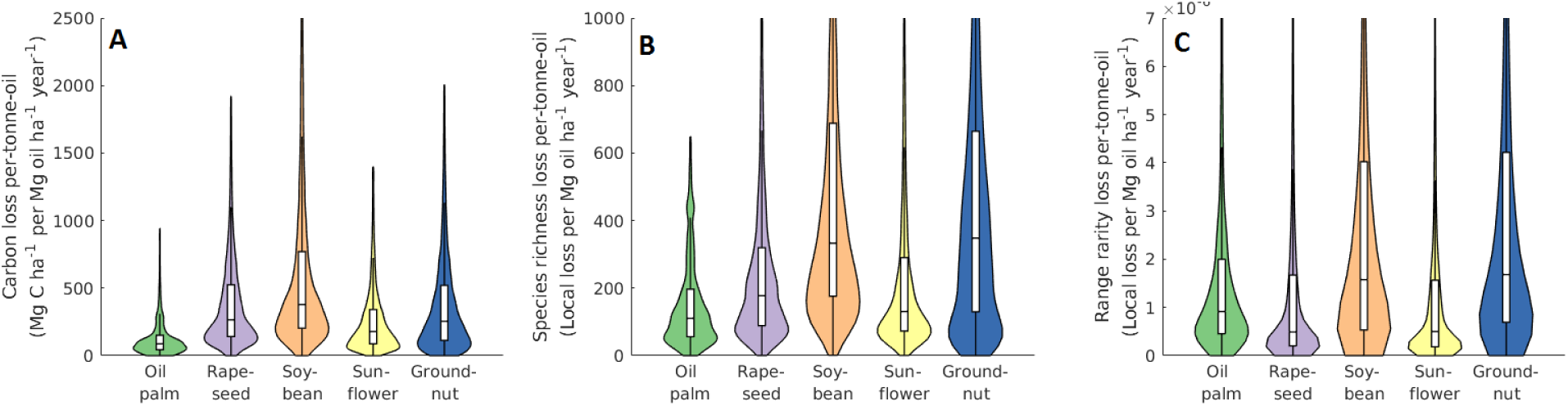
Crop-specific distributions of (A) carbon loss, (B) species richness loss and (C) range rarity loss per-tonne-oil on global harvested areas in 2010. Boxes show medians and upper and lower quartiles.

Following on from our analysis of current growing areas, we investigated how future global demand for vegetable oil could be met with the lowest possible additional environmental impact. Closing yield gaps (i.e. the difference between actual and agro-climatically attainable yields) on current croplands could substantially increase vegetable oil production with minimal additional environmental impact; however, realising this is not trivial, and demand levels in coming decades are likely to rise even beyond theoretically achievable production on existing growing areas ^21^. This requires us to identify where future crop expansion will cause the least environmental damage. We therefore repeated our analysis using projected yields on agro-climatically suitable areas (Fig. S2), which represent potential future growing locations (Methods). There are a number of geographical areas where predicted impacts per-tonne-oil are substantially lower than typical values on current croplands (Fig. 2). Specifically, carbon loss per-tonne-oil is predicted to be lowest for oil palm in areas of tropical South America and Central Africa, and for groundnut planted in Central-East Asia (Fig. 2A,B). Species richness loss per-tonne-oil is estimated to be lowest if areas are planted with oil palm in parts of Southeast Asia and Central Africa, and rapeseed and sunflower in Central and Eastern Europe (Fig. 2C,D). Range rarity loss per-tonne-oil is predicted to be lowest for rapeseed and sunflower planted in Central and Eastern Europe and parts of central North America, and for groundnut planted in East China (Fig. 2E,F). We highlight that even within relatively small geographical regions (e.g. within Borneo, or within Sumatra), there can be considerable spatial heterogeneity in the predicted environmental impacts per-tonne-oil, of up to an order of magnitude. We also note that due to the limited spatial resolution of the data (∼10km at the equator), these predictions do not replace the need for detailed on-the-ground yield and impact assessments prior to land use change.

**Fig. 2:**
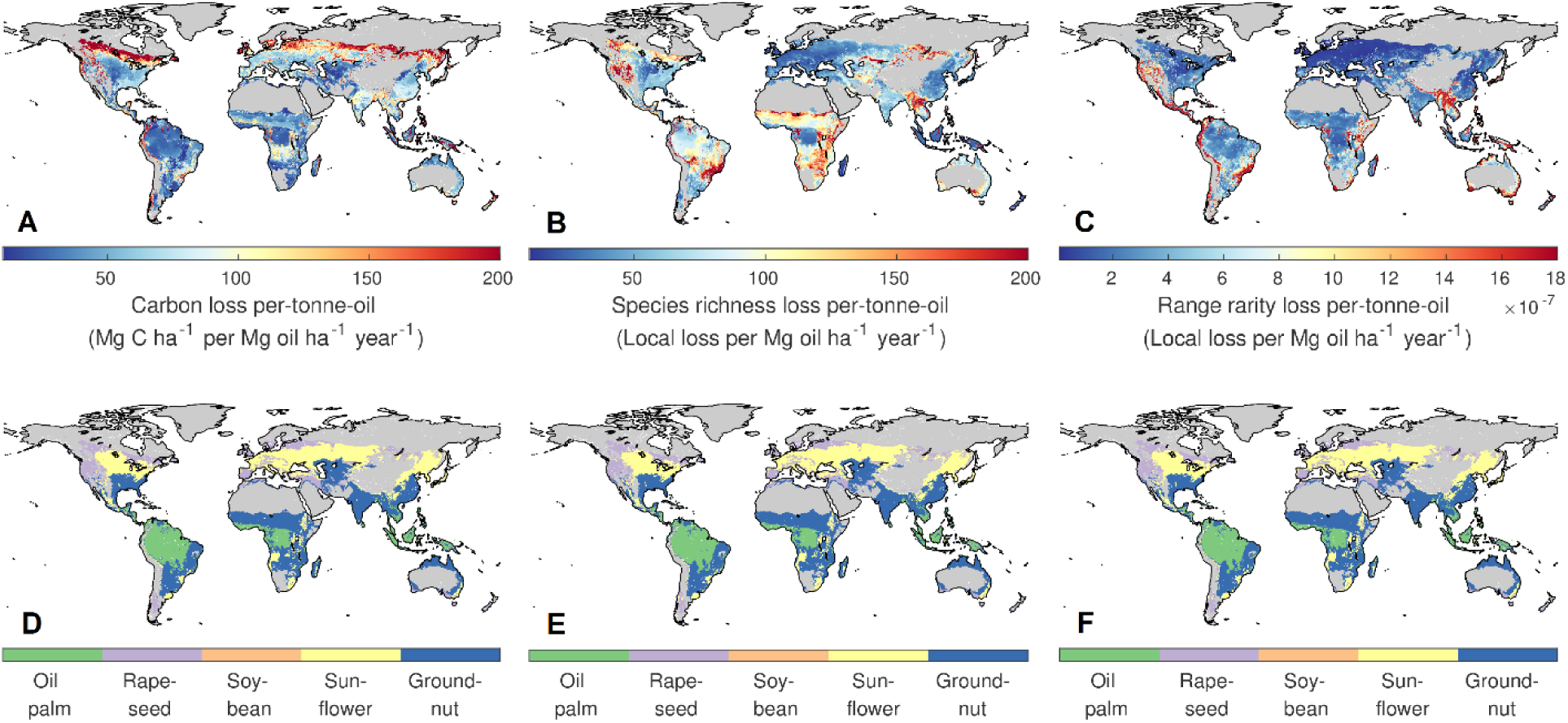
Maps of per-tonne-oil (A) carbon loss, (B) species richness loss (C) range rarity loss, when selecting oil crops that locally minimise these impacts (D,E,F, respectively). Coloured areas represent all agro-climatically suitable locations for oil crops (of which only a tiny fraction needs to be cultivated to meet current and future global vegetable oil demand). More than one crop may be able to grow in a coloured location, but only the choice shown in (D–F) will minimise local per-tonne-oil impacts to the lowest possible values, shown in (A–C). Thus, vegetable oil is produced at the lowest possible impacts when dark blue areas in (A–C) are cultivated with the appropriate crop shown in (D–F). Impact calculations are based on biodiversity and carbon stocks of the natural habitat. Fig. S3 provides a high-resolution version of this figure.

We estimate that the carbon and biodiversity impacts of closing current yield gaps and expanding future croplands into optimal areas would be substantially lower than under a business-as-usual scenario of increasing current production. We modelled the environmental impacts of a gradual expansion of oil crops into areas potentially available for future agricultural use (Methods), starting with areas where per-tonne-oil impact is lowest (i.e. dark blue areas in Fig. 2A–C). This simulates the minimal achievable impact of each crop for varying levels of global vegetable oil production. We note that this approach only considers agro-ecological parameters, and does not account for socio-economic factors such as local infrastructure, the cost of land and labour, distance to markets, and policies. We find that this strategy would drastically reduce carbon, species richness and range rarity impacts estimated to occur under a business-as-usual expansion of oil croplands to meet a demand of 340 million tonnes projected for 2050 (Fig. 3). This is in line with previous research highlighting the importance of spatial planning for minimising the environmental impacts of agricultural expansion ^22^. Our results suggest that oil palm outperforms its alternatives with regard to minimising carbon loss and species richness loss per-tonne-oil, if new growing areas were established in optimal locations, while sunflower and rapeseed rank higher with regard to range rarity loss (similar to our findings for current growing areas; Fig. 1). Trade-offs between impact measures are very small, with only marginally higher carbon and biodiversity losses when impacts are minimised simultaneously (Fig. S4).

**Fig. 3:**
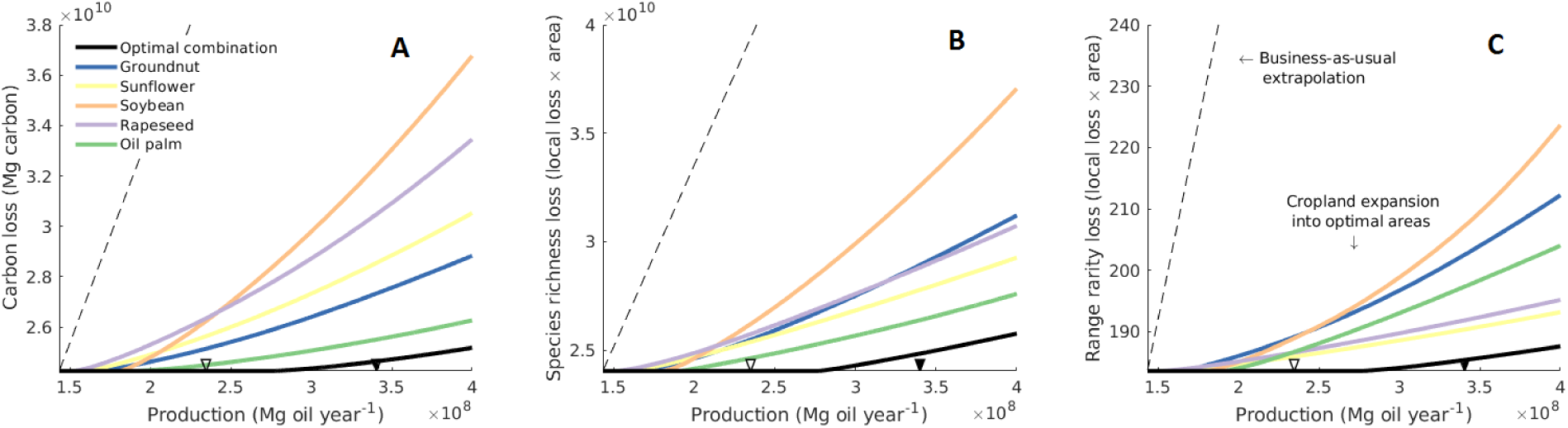
Minimal achievable impacts of future vegetable oil production in terms of (A) carbon loss, (B) species richness loss and (C) range rarity loss, based on closing yields gaps and optimally expanding 2010 cropping areas. Coloured lines represent yield gap closure (which does not increase impacts) and the subsequent optimal expansion of individual crops into currently unused land (Methods), ordered by increasing impact per-tonne-oil score. Black lines represent optimal combinations of crops. Dashed lines correspond to a business-as-usual scenario based on a linear extrapolation of 2010 production and impact data. Coordinate origins correspond to production and impacts in 2010. White and black triangles represents projected global demand for 2027 ^10^ and an upper estimate for 2050 ^11^, respectively.

The differences between oil crops in Fig. 3 are small compared to the effect of planting any of the individual crops in optimal locations. This matches our results for current growing areas, showing that growing location generally affects the environmental impact of vegetable oil production more than crop type. Economic, infrastructural and regulatory policies and strategies aimed at steering vegetable oil production towards optimal areas ^23^ are therefore likely to be much more effective at reducing impacts than categorically substituting one oil type for another.

Prioritising crop expansion into agriculturally degraded land has been suggested as a means to minimise environmental impacts ^2,16,24,25^. Whilst the short-term damage associated with this strategy is much lower than in the case of natural habitat conversion, it is important to note that establishing cropland on degraded areas can prevent the regeneration of biodiversity and carbon stocks that may occur otherwise. Carbon stocks and species richness can reach pre-disturbance levels within a century, though the restoration of community composition can take much longer ^26–32^. In some cases, the long-term environmental impacts of planting oil crops in areas of degraded versus old-growth ecosystems could therefore in theory be similar. However, given the current lack of effective policies that could practically ensure the long-term regeneration of degraded lands ^33^, the uncertainty in predicting local recovery success ^32^, and the importance of primary habitats as biodiversity sources ^34^, we echo calls to prioritise future oil crop expansion on degraded areas, specifically those that are located ^25^ within optimal (i.e. dark blue) areas in Fig. 2A–C.

In our analysis we have not accounted for potential economic feedbacks related to shifts in the proportion of global oil demand supplied by individual crops. For example, it has been suggested that a relative increase in palm oil production, leading to a lower price of vegetable oil, may ultimately increase total vegetable oil demand and thereby incentivise further land conversion ^13,35^. However, such rebound effects are not inevitable; policy options that can allow them to be overcome include zoning land for oil crop production and for conservation, creating economic incentives such as land taxes and subsidies, and promoting robust certification standards ^23,36^.

Whilst carbon and biodiversity loss are arguably the two most controversial environmental impacts associated with vegetable oil production, there are other impacts that we have not included in our analysis due to a lack of suitable spatial data. Land clearing, irrigation practices, fertiliser and pesticide use can reduce air and water quality, increase soil erosion and negatively affect a range of ecosystem functions ^37^. Localised estimates of other impacts are often only available for agricultural land use in general rather than specific oil crops ^37^, hindering comparisons between them. Global data to allow the quantification of such impacts, and the trade-offs between them, is therefore urgently needed. Our analysis also does not account for socio-economic impacts of vegetable oil production. Oil palm expansion, in particular, has been associated with cases of land use conflicts with local residents ^38^. At the same time, palm oil production has also contributed to increasing employment and improving social services ^39^, although these effects are spatially heterogeneous ^40^. Strong socio-economic standards are needed to ensure land rights, acceptable labour conditions, and the equitable distribution of benefits among companies, government, and locals ^40^.

Our findings demonstrate that the environmental impacts of oil palm are complex. Per-tonne-oil impacts on carbon and species richness loss are lower than for four other major oil crops, while impacts on small-ranged species rank third after sunflower and rapeseed. These results challenge earlier research and public perceptions of oil palm, which generally do not take into account the area requirements of alternative crops for producing oil. Our analyses provide vital quantitative evidence when considering the potential impacts of switching production from oil palm to other crops ^16^, adding important nuance to ongoing debate. We have only estimated impacts on three environmental metrics; further research on other impacts of oil production such as those on ecosystem services and socio-economic conditions is vital to inform decision-making. Significantly, we have shown that the location of oil crops plays a crucial role in determining their impacts. As such, if vegetable oil demand increases as predicted, closing yield gaps and prioritising the expansion of croplands in areas with the lowest predicted environmental costs per-tonne-oil will likely help reduce negative impacts much more than substituting one oil type for another. Such strategies need to be coupled with efforts to reduce overconsumption and waste ^41^ of vegetable oils, thereby minimising their global environmental footprint.

## Acknowledgements

The authors are grateful to Juliet Vickery, Andrew Callender, Alice Ward-Francis, Genevieve Hayes, Heather Ducharme, Corli Pretorius, Kimberly Carlson, Mario Herrero, Ben Phalan, Andrew Balmford and Andrea Manica for comments. This work was supported by the Cambridge Conservation Initiative Collaborative Fund (CCI-06-16-008) and the Arcadia Fund.

## Author contributions

R.M.B, A.P.D., C.T., S.E.B., D.C., P.F.D. and F.J.S. designed the study. R.M.B. conducted the analysis and wrote the initial draft. All authors interpreted the results and revised the manuscript.

## Methods

### Current and potential growing areas and yields

We used 5 arc-minute (∼10 km at the equator) global maps of harvested areas and yields for oil palm, soybean, rapeseed, sunflower and groundnut for the year 2010 ^17,42^, the most recent spatial crop data. To assess the robustness of the results shown in Fig. 1 with respect to the underlying crop data, we additionally repeated the analysis using an independent dataset available for the year 2000 ^43^, and obtained qualitatively identical results in terms of the ranking of crops for the different environmental impacts. Conversion factors ^44^ were used to derive oil yields from crop yields. As we focus on vegetable oil production here, we do not consider possible by-products of oil crop cultivation such as oil cake. For each crop, we used potential growing areas and agro-climatically attainable oil yield (used in Fig. 2 and 3) assuming rain-fed water supply and high input management ^45^ (Fig. S3). These potential yields were also used to estimate crop-specific yield gaps on current growing areas (used in Fig. 3). We note that these data do not account for new crop varieties; for example, potential palm oil yields may reach 10 Mg ha^-1^ year^-1 13,15,37,46^, which represents an increase of up to 27% compared to the highest agro-climatically attainable yield levels used here. The land that is available for potential cropland expansion (used in Fig. 3) in a given grid cell was defined as the fraction that is not currently taken up by cropland, pasture and urban areas, for which we used a recent 5 arc-minute global dataset, representative of the year 2016 ^47^.

### Carbon loss

Following ref. ^18^, we estimated the local carbon impact associated with the conversion of natural land cover to cropland in any given 5 arc-minute grid cell as the difference between (i) local carbon stocks in potential natural vegetation and soils (Fig. S5A), and (ii) local carbon stocks under agricultural land cover. The change in vegetation carbon from land conversion was calculated as the difference of carbon stocks in potential natural vegetation ^18^ and in crops ^24,43^. The loss of soil organic carbon (SOC) from land conversion to cropland – following ref. ^18^ and supported by empirical meta-analyses ^48–52^ – was estimated as 25% of potential natural SOC, where we used a recent estimate ^53^ for the latter. On peatland ^54^, we assumed a 100% loss of potential SOC in the course of the depletion of SOC from peat drainage. This conservative assumption results in disproportionally higher SOC losses for oil palm, the only crop out of the five considered for which a non-negligible portion is currently grown in areas with high peat occurrence. It is also likely an overestimation given that significantly lower SOC loss percentages on peatland can be achieved by means of appropriate management practices ^55^.

In addition to emissions associated with carbon losses from the replacement of natural vegetation and from soil conversion, considered here, nitrous oxide emissions from fertiliser use represent an additional source of agricultural greenhouse gas emissions ^54^ that we have not included in our calculation. Whilst a spatial dataset of global fertiliser application rates has been derived for the year 2000 ^56^, based on which emissions can be estimated ^54^, extrapolating these data to the 2010 cropland data, and to agro-climatically suitable potential future growing areas considered in our analysis would involve high uncertainty, as the relationships between fertiliser use and local environmental and economic factors are not trivial ^56^. That said, we find that the ranking of oil crops with respect to average nitrous oxide emissions per-tonne-oil on growing areas in 2000 (Fig. S6) is indeed identical to the ranking for average carbon losses (Fig. 1). In particular, average nitrous oxide emissions per-tonne-oil are lower for oil palm than for the other four crops. Furthermore, Fig. 1 and Fig. S6 demonstrate that nitrous oxide emissions, even when aggregated over long time periods, are small when compared to emissions associated with land use change (both after conversion to CO2eq) considered in our analyses, and are therefore very unlikely to affect our conclusions.

### Biodiversity loss

Following ref. ^19^, we estimated the local biodiversity impact associated with the conversion of natural land cover to cropland in any given 5 arc-minute grid cell as the difference between (i) local biodiversity under potential natural vegetation, and (ii) local biodiversity under agricultural land cover, as follows. We used worldwide spatial extents of occurrence (EOO) for all known birds ^57^, mammals and amphibians ^58^. Using species-specific altitudinal ranges and habitat preferences, we refined the EOO according to local elevation and given land cover: A species was considered present in a 5 arc-minute grid cell under potential natural land cover if (i) its EOO contained the grid cell centre, (ii) the local elevation ^59^ was contained in the species’ altitudinal range, and (iii) the species’ list of suitable habitats included the local potential natural vegetation ^60^ according to the matching in Table S2 (Fig. 5B). Analogously, a species was considered present in an oil palm plantation, or in a rapeseed, soybean, sunflower or groundnut field in a given grid cell, if conditions (i) and (ii) were met, and if the species’ list of suitable habitat types includes the IUCN habitat categories ‘Plantations’ or ‘Arable Land’, respectively. Combining the resulting two maps provides an estimate of the species lost in any given 5 arc-minute grid cell when potential natural vegetation is converted to the relevant cropland type.

Based on these data, we used two alternative metrics to assess grid cell-specific biodiversity loss due to land conversion. First, we used the decrease in absolute species richness, i.e. the (unweighted) sum of the estimated local loss of bird, mammal and amphibian species. Second, we used the inverse species-specific potential natural range (i.e. the area of the refined EOO for the potential natural land cover) as weights in the above sum. In this way, rare species, characterised by small geographical ranges, carry more weight than geographically widespread species (Fig. 5C). This metric – range rarity – has been advocated as a biodiversity measure alternative to species richness ^20,61–63^.

The species habitat data do not contain further differentiation within the IUCN habitat categories ‘Arable land’ and ‘Plantation’. Our assessments of the loss of species richness due to land conversion are thus potentially an underestimation, given that, in general, fewer species are able to live, for example, in a rapeseed field, than those listed as tolerating ‘Arable land’ overall. The available data do not allow us to consider variation in the mitigation of local loss of species due to land conversion by means of more biodiversity-friendly cropland design. Whilst some studies have advocated such strategies ^37,64–66^, strong evidence has been presented against their effectiveness ^3,67–71^.

### Environmental impacts per-tonne-oil

Using the above data, we calculated the local environmental impact per-tonne-oil, i.e. the ratio of local carbon or biodiversity impact to local oil yield, for each crop and 5 arc-minute grid cell. The distributions of these values across current harvested areas are shown in Fig. 1. Grid-cell-specific impacts per-tonne-oil in Fig. 2 are calculated in an analogous way (using the fact that our estimation of carbon and biodiversity losses from land conversion can be applied on both existing croplands and currently uncultivated areas), but use potential yields on agro-climatically suitable areas (Fig. S3) instead of current yields.

## Supplementary Material

**Fig. S1:**
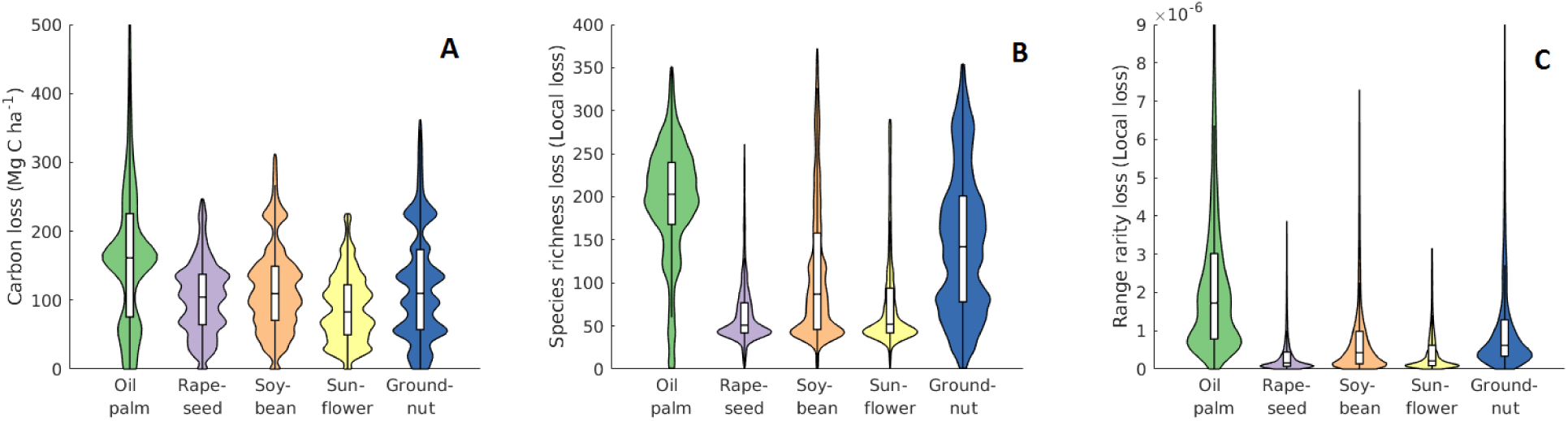
Crop-specific distributions of absolute (i.e. not relative to oil yields) (A) carbon loss, (B) species richness loss and (C) range rarity loss on global harvested areas in 2010. Boxes show medians and upper and lower quartiles.

**Fig. S2:**
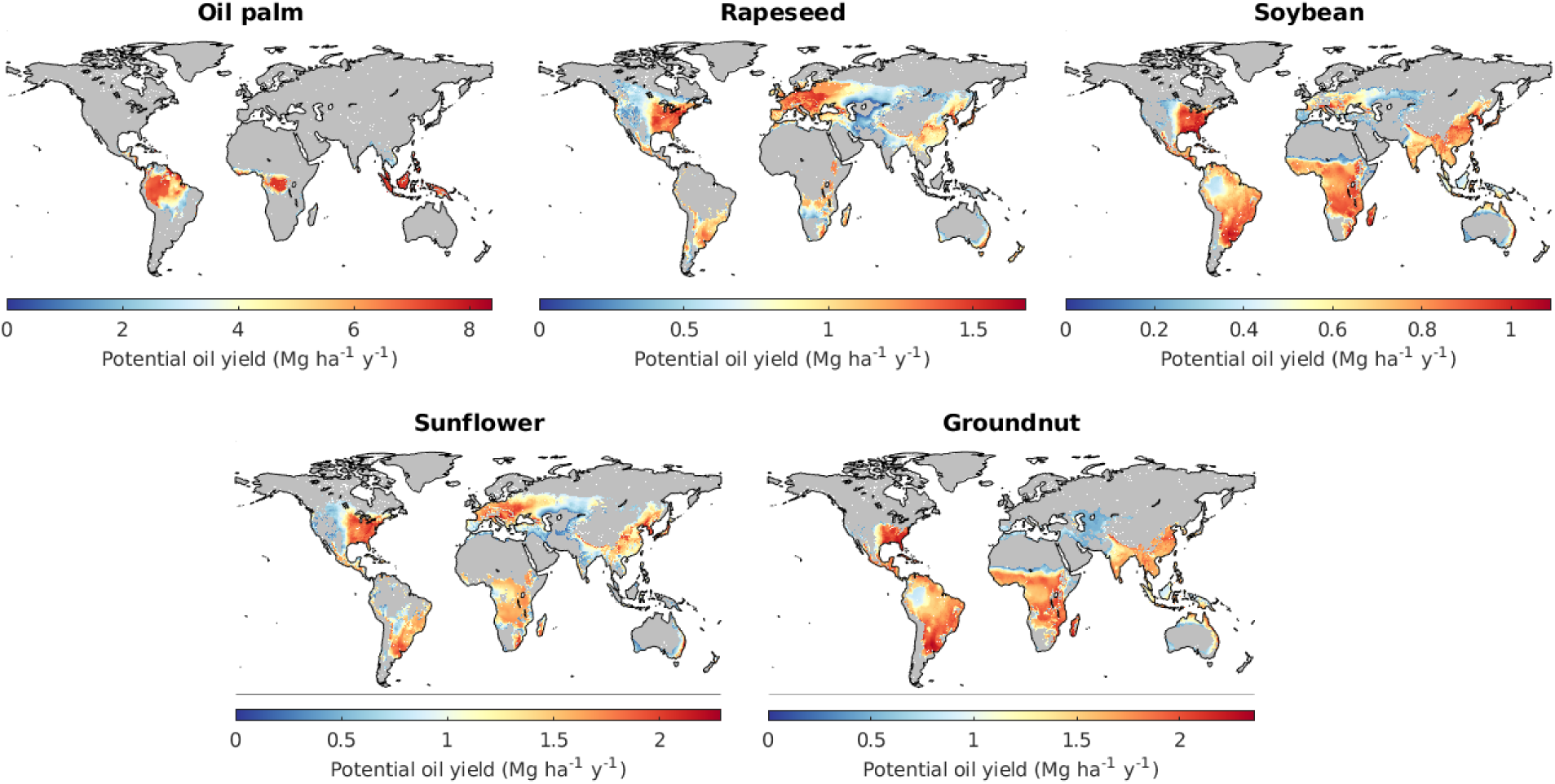
Crop-specific agro-climatically suitable growing areas and potential yields, assuming rain-fed water supply and high input management ^45^. Note the different scales of the colour bars. For each crop, the area required to produce one tonne of vegetable oil in a specific grid cell is given by the inverse of the local yield shown.

**Fig. S3:**
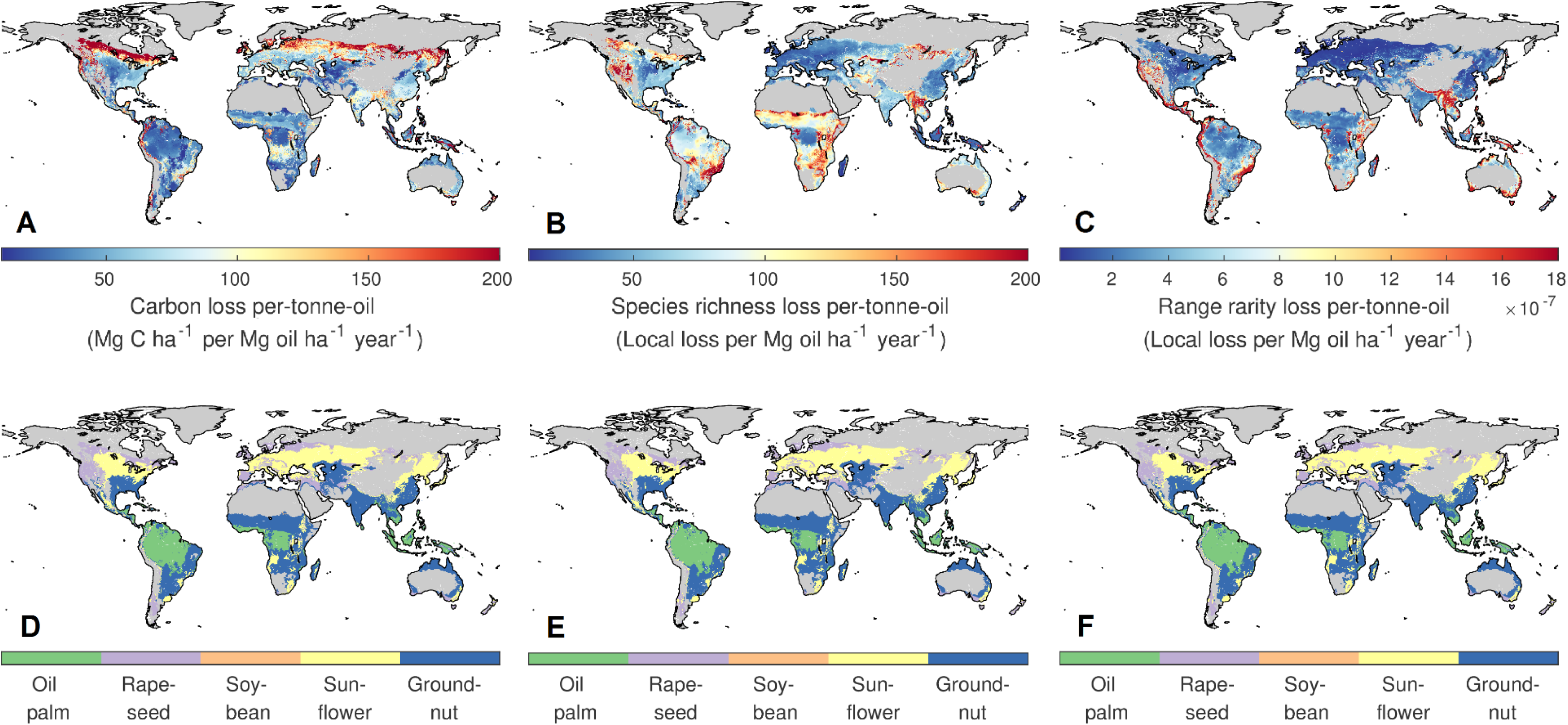
High-resolution version of Fig. 2.

**Fig. S4:**
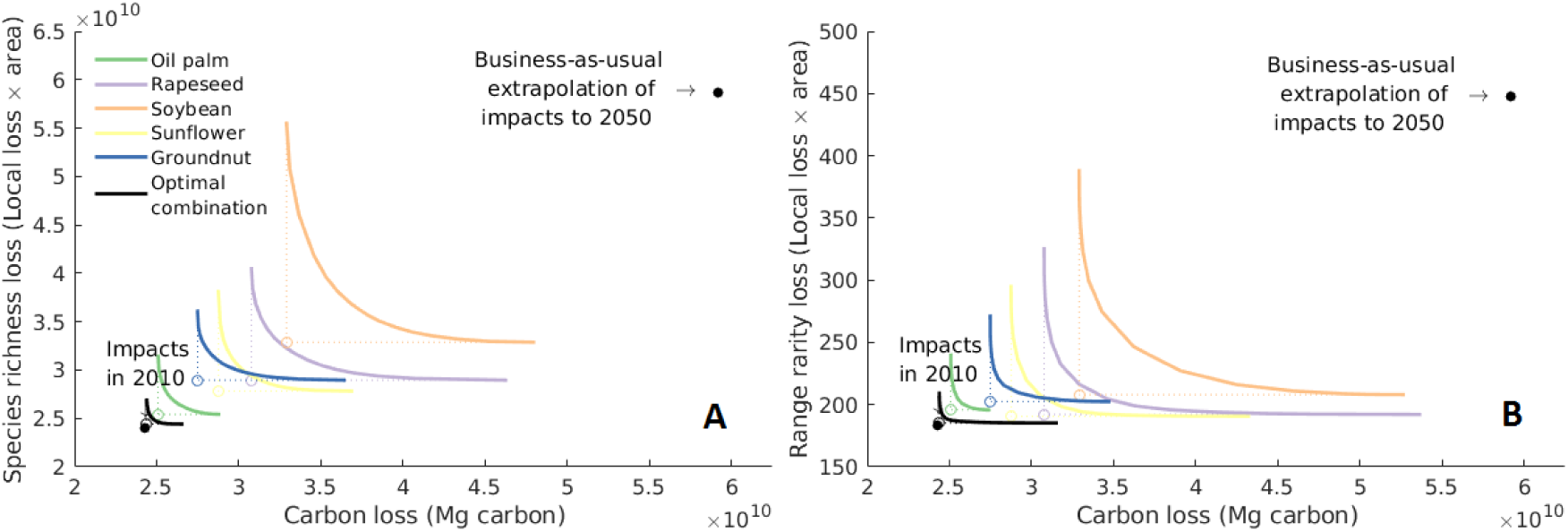
Trade-offs between reducing carbon and biodiversity loss. All lines correspond to a production level of 340 million tons of oil estimated for 2050 ^11^. For each crop (coloured lines) and an optimal combination of crops (black line), lines represent the minimum simultaneously achievable carbon and biodiversity impacts when yield gaps are closed and growing areas are optimally expanded. Open coloured circles correspond to the hypothetical scenario of no trade-offs. The high convexity of the lines demonstrates that trade-offs are small.

**Fig. S5:**
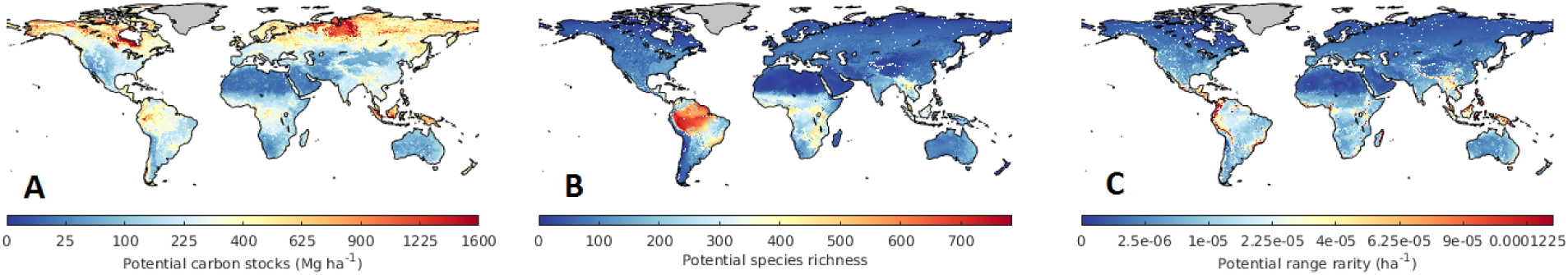
Potential natural (A) carbon stocks (vegetation and soil organic carbon), (B) species richness and (C) range rarity.

**Fig. S6:**
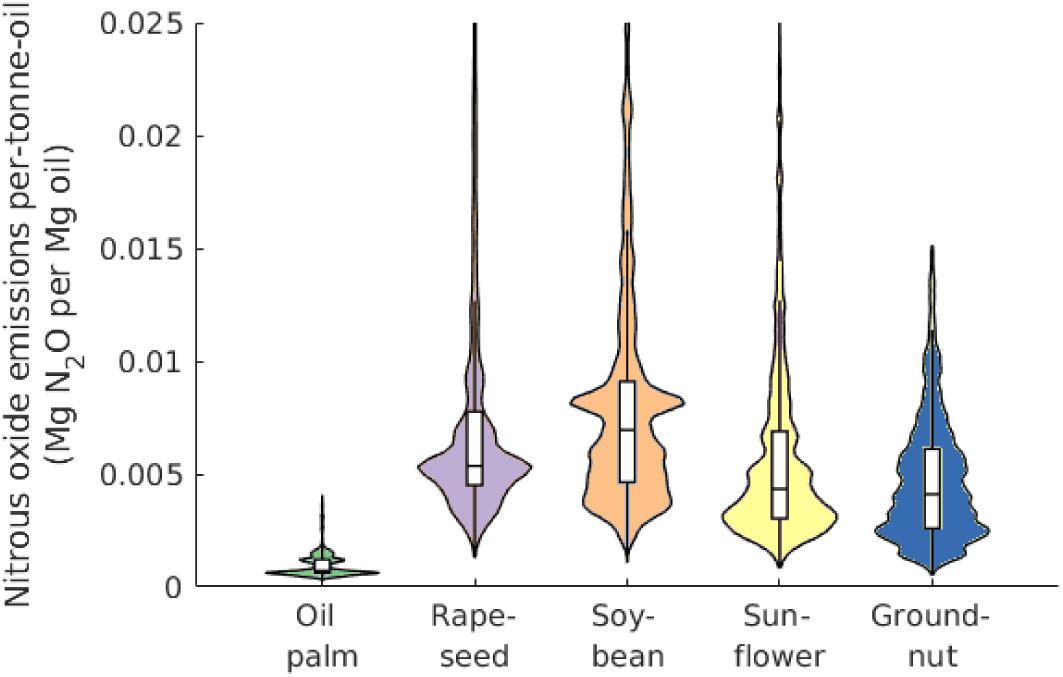
Crop-specific distributions of nitrous oxide emission per-tonne-oil on growing areas in 2000. Direct and indirect nitrous oxide emissions were calculated based on a spatial dataset of nitrogen application rates for the year 2000 ^56^, using a standard methodology ^54^. For consistency, the calculation of N_2_O emissions per-tonne-yield is based on yields and harvested areas derived for the year 2000 ^60^, which has been used in the generation of the fertiliser data. Boxes show medians and upper and lower quartiles.

**Table S1:** Matching of potential natural land cover categories used by Ramankutty and Foley ^60^ (R&F) and IUCN categories for species habitats.

| IUCN species habitat categories | Equivalent R&F potential natural land cover categories |
| --- | --- |
| Desert | Desert<br>Polar desert / Rock / Ice |
| Forest – Boreal | Boreal Evergreen Forest / Woodland<br>Boreal Deciduous Forest / Woodland<br>Mixed Forest <sup>(1)</sup> |
| Forest – Subarctic / Subantarctic | Boreal Evergreen Forest / Woodland<br>Boreal Deciduous Forest / Woodland<br>Mixed Forest <sup>(1)</sup> |
| Forest – Subtropical / Tropical | Tropical Evergreen Forest / Woodland<br>Tropical Deciduous Forest / Woodland<br>Mixed Forest <sup>(1)</sup> |
| Forest – Temperate | Temperate Broadleaf Evergreen Forest / Woodland<br>Temperate Needleleaf Evergreen Forest / Woodland<br>Temperate Deciduous Forest / Woodland<br>Mixed Forest <sup>(1)</sup> |
| Grassland | Savanna <sup>(2)</sup><br>Grassland / Steppe<br>Tundra |
| Marine | Water Bodies (Oceans) |
| Rocky areas | Polar desert / Rock / Ice |
| Savanna | Savanna |
| Shrubland | Dense Shrubland<br>Open Shrubland |
| Wetlands <sup>(3)</sup> | Grassland / Steppe<br>Dense Shrubland<br>Open Shrubland |
|  | Cropland categories |
| Arable Land | Rapeseed, soybean, sunflower or groundnut field |
| Plantations | Oil palm plantation |

<sup>(1)</sup> 'Mixed forests' do not have a direct equivalent in the IUCN categories. We defined a species as potentially present in a grid cell where the potential natural vegetation is 'Mixed forests' if its habitat categories included any type of forest. A different approach is unlikely to affect our results significantly as the number of such grid cells is very small.
<sup>(2)</sup> The definition of 'Savanna' in the IUCN categories is much more narrow than the one used by R&F, who also use the term for what would be considered 'Grassland' in the IUCN categories. <sup>(3)</sup> The categories used by R&F do not include inland wetlands. A species whose habitats include wetlands, and whose other habitat categories do not include the potential land cover, was listed as potentially present in a grid cell if the potential land cover is grassland or shrubland, and if the grid cell contains wetlands <sup>72</sup>.

